# A single-sample circadian biomarker that performs across populations and platforms

**DOI:** 10.1101/820811

**Authors:** Gang Wu, Marc D. Ruben, Lauren J. Francey, David F. Smith, Joseph D. Sherrill, John E. Oblong, Kevin J. Mills, John B. Hogenesch

## Abstract

**Background:** For circadian medicine to influence health, such as when to take a drug or undergo a procedure, a practical way to measure body time is needed. Recent machine learning algorithms show that gene expression data from blood and skin can provide reliable estimates of body time. However, for clinical viability, a biomarker must be easily measured and generalizable to a broad population. It is not clear that any circadian biomarker yet satisfies these criteria.

**Results:** We analyzed 24 h molecular rhythms in human dermis and epidermis at three distinct body sites, leveraging both longitudinal and population data. Circadian clock function was strongest in the epidermis, regardless of body site. We identified a 12-gene biomarker set that reported circadian phase to within 3 hours from a single sample of epidermis—the skin’s most superficial layer. This set performed well across body sites, ages, sexes, and detection platforms.

**Conclusions:** This research shows that the clock in epidermis is more robust than dermis regardless of body site. To encourage ongoing validation of this biomarker in diverse populations, diseases, and experimental designs, we developed SkinPhaser—a user-friendly app to test biomarker performance in datasets (https://github.com/gangwug/SkinPhaser).

## Background

In the last 50 years, dozens of clinical studies showed that dosing time-of-day can impact the efficacy and safety of many different types of medical treatments [1,2]. We know from multi-tissue studies in mice [3] and humans [4] that thousands of rhythmically expressed genes encode known drug targets. Circadian medicine aims to incorporate this knowledge to improve treatment outcomes. This requires a reliable measure of a patient’s internal time as wall time does not necessarily equate to body time. Accumulating evidence shows interpersonal variation in the timing of physiology and behavior due to genetics, age, sex, and lifestyle [5–8]. Markers of body time are therefore in high demand.

Previous research focused on dim-light melatonin-onset (DLMO) as a measure of body time [9,10]. However, DLMO is inconvenient, costly, difficult to standardize, and thus rarely used clinically. With the development of high-throughput molecular detection platforms and computational techniques, recent efforts shifted to machine learning predictions based on the transcriptome or metabolome from one or two samples of whole blood [11–16], or a specific blood cell type (e.g., CD14^+^ monocytes) [17]. Using these developed machine learning approaches, the DLMO phase (or sampling time) could be accurately assigned within 3 h from a single blood sample. However, these studies were small (≤ 74 subjects) and limited to younger subjects (18-41 years of age). Developing and validating circadian biomarkers in the broader population remains a major challenge.

Previously, we developed a 29-gene biomarker set from ~300 human forearm skin samples. From a single biopsy, this set reliably estimated circadian phase to within 3 h of expected time [18]. However, for clinical use, critical questions remain. Does this biomarker set perform across different body areas? In different populations? On different platforms? With different experimental designs?

Using an experimental design that captures the advantages of both longitudinal and population-based studies, we analyzed 24 h molecular rhythms in human dermis and epidermis at three distinct body sites. Subjects in the longitudinal group (n=20, aged 21-49 y) each donated one skin punch biopsy every 6 h across a day. Subjects in the population group (n=154, aged 20-74 y) each donated one punch biopsy from forearm, buttock, and cheek without recording time. For all biopsies, the dermis was separated from the epidermis by laser capture microdissection (LCM) and then profiled on gene expression arrays. We applied CYCLOPS [19] to order the population samples where biopsy time was not recorded.

We found that circadian clock function was strongest in the epidermis, regardless of body site. Based on this, we applied ZeitZeiger [20] and identified a 12-gene biomarker set from a single epidermal sample that reported circadian phase to within 3 h. This set performed well across body sites, ages, sexes, and detection platforms and represents a forward path to clinical application of circadian medicine.

## Results

### A functional clock in the human dermis

To explore molecular rhythms in human dermis, forearm skin samples were collected from the longitudinal group every 6 h from 20 individuals (subject 115 had one missing time point and was excluded; Additional file 1: Table S1). The dermal layer was separated from the epidermal layer using LCM. Using MetaCycle [21], we identified 182 circadian genes (*P* < 0.05; Additional file 1: Fig. S1A) in the dermis. The distribution was bimodal (Additional file 1: Figure S1B) with peak phases clustered at 8-9 AM and 8-9 PM, as previously seen in epidermis [18]. Morning-time genes were enriched for immune-related and G protein-coupled receptor (GPCR) signaling, whereas the evening was marked by genes involved in transcription, lipid and lipoprotein metabolism, and development (Additional file 1: Figure S1C and S1D) (Phase Set Enrichment Analysis (PSEA) [22]).

### Clock strength varies between skin layers

We next compared clock gene oscillations between forearm dermis and epidermis. The peak phases of expression for six clock genes (*ARNTL*, *NR1D2*, *HLF*, *PER3*, *PER2* and *NFIL3*; *P* < 0.1) (Fig. 1) between skin layers were generally consistent in subjects with the same circadian phase. For example, subjects 116 and 119 were phase delayed in both skin layers compared to all other subjects in terms of both gene expression (Fig. 1) and salivary cortisol levels (Additional file 1: Figure S1E). Although clock gene phases were similar between skin layers, the strength of their oscillations were not. For each of the six clock genes that were rhythmic in both layers, amplitudes were higher in the epidermis. For example, the median relative amplitude (rAMP) of *PER3* was two-fold greater in the epidermis compared to dermis (0.59 vs. 0.29) (Additional file 1: Figure S1F).

**Fig. 1.**
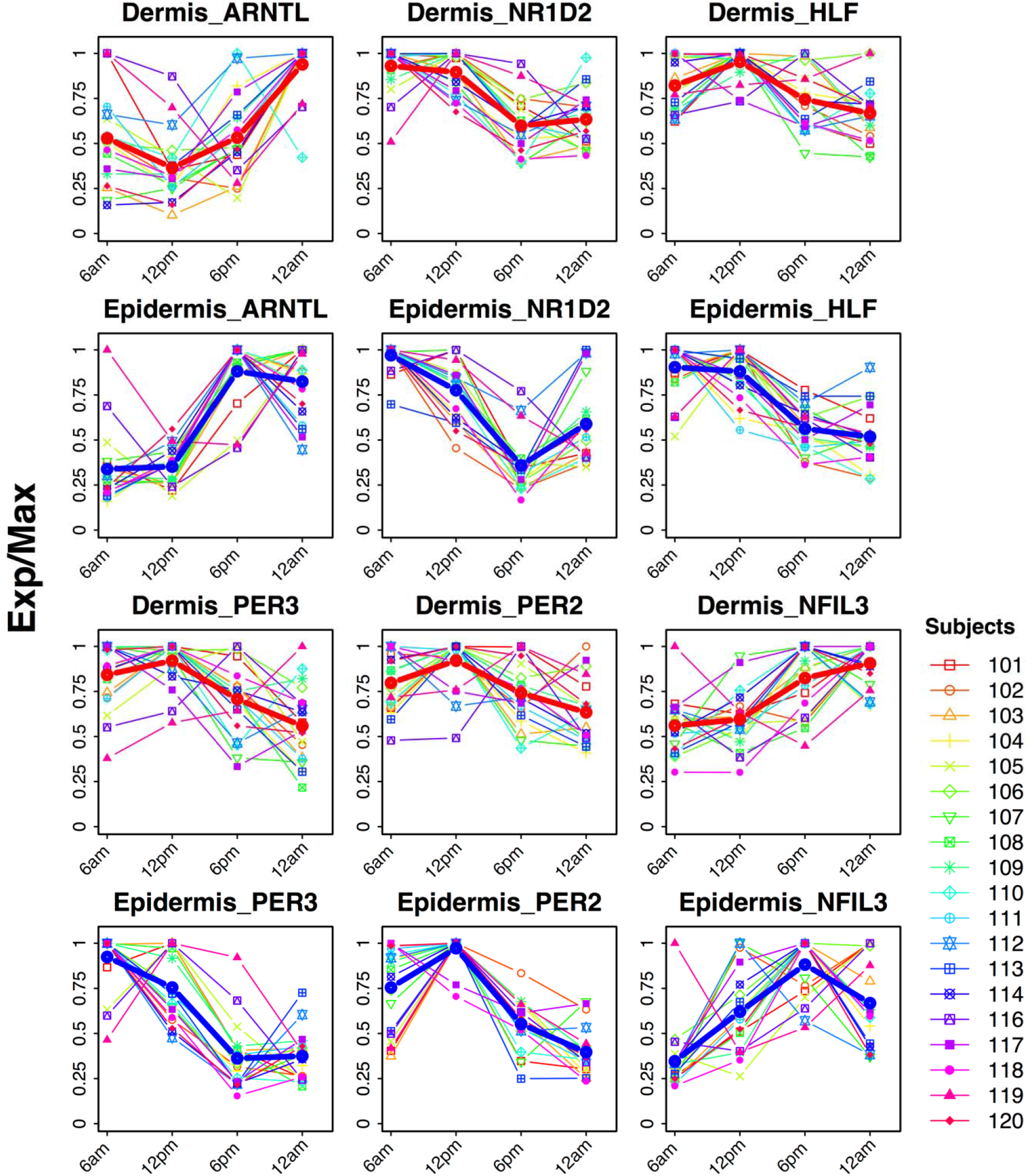
Expression profiles of overlapped clock and clock-associated genes in human dermis and epidermis. Forearm punch biopsies were collected every 6 h for 1 day starting at 12 PM. Expression profiles from 19 subjects over time are shown for dermis (rows 1 and 3) and epidermis (rows 2 and 4). Red and blue lines represent the average expression profile among all subjects for dermis and epidermis, respectively. Exp/Max indicates the expression value at each time point normalized to the maximum expression across four time points.

To evaluate clock function in the broader population, we next analyzed clock gene expression from 154 subjects, aged 20-74 y. Each subject contributed three punch biopsies from cheek, forearm, and buttock [23]; sampling time was not recorded. To compare robustness of clock oscillations between different skin layers and body sites, we computed correlation matrices of clock and clock-associated genes [24] for each of the six conditions (e.g., epidermis-cheek, dermis-cheek, etc.). Each of these matrices were compared to a reference matrix constructed from 12 mouse tissues (Additional file 2). To quantify this, a Mantel test Z-statistic was computed, which provided a measure of similarity between the mouse reference and each of the six conditions. Larger values indicate stronger clock oscillation. As expected, forearm epidermis (Z-statistic = 33.3; Fig. 2A) was stronger than dermis (Z-statistic = 29.6; Fig. 2B). This was also true in the non-sun-exposed buttock, with a larger Z-statistic value in epidermis (Z-statistic = 34.7; Fig. 2C) than dermis (Z-statistic = 32.1; Fig. 2D). There was no obvious difference in clock robustness between cheek epidermis (Z-statistic = 35.3; Fig. 2E) and dermis (Z-statistic = 35.6; Fig. 2F). When combining all longitudinal and population study samples from three body sites, the epidermis (Z-statistic = 37.6; Fig. 3A) was stronger than dermis (Z-statistic = 32.5; Fig. 3B). Of note, although the dermal clock is weaker, it is still functional (Mantel test *P* < 5.0e−06).

**Fig. 2.**
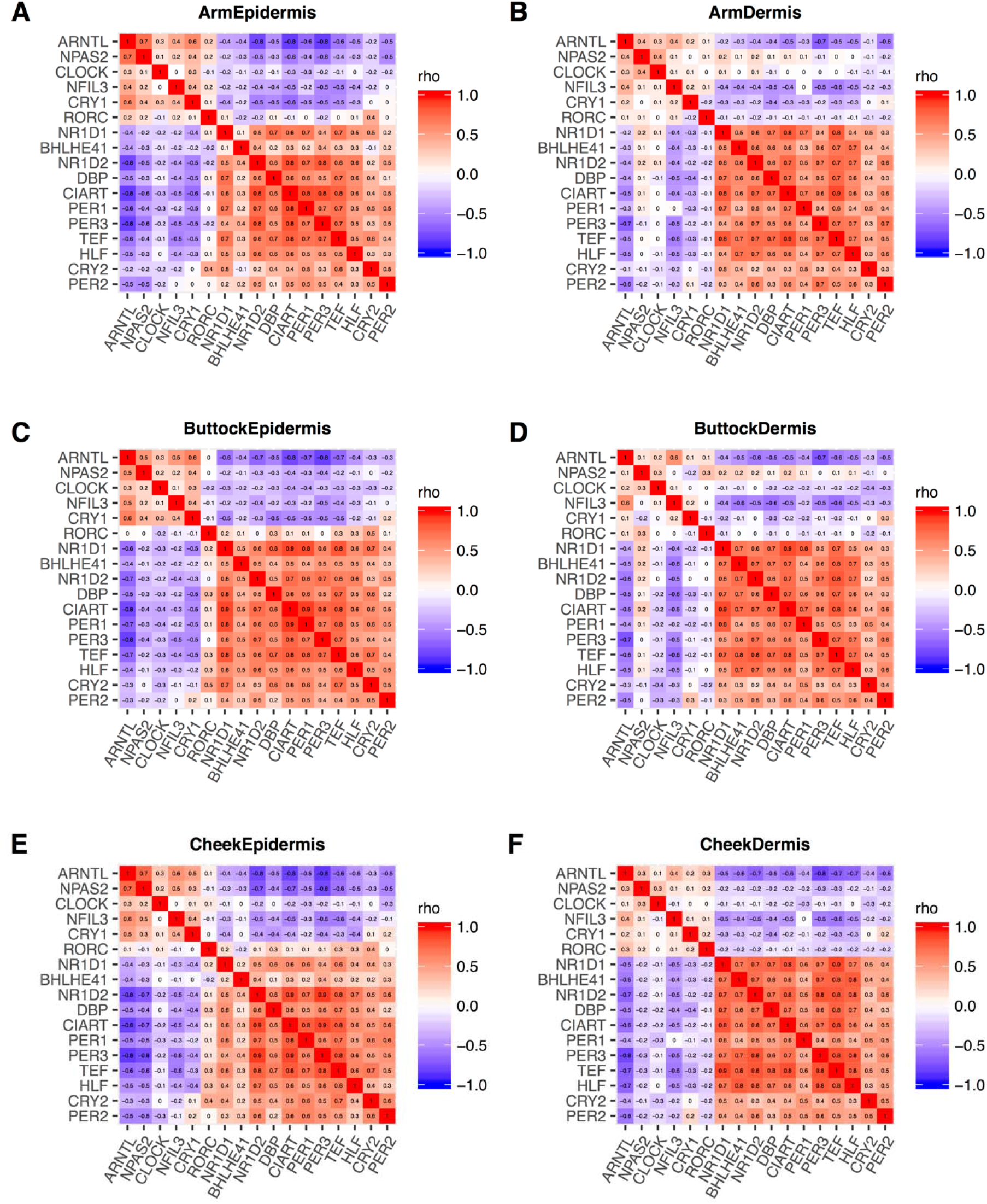
Evaluation of circadian clock robustness in epidermis and dermis across different body sites. The heatmaps of Spearman’s rho correlation values for clock and clock-associated genes [24] were drawn for epidermis (A, C, E) and dermis (B, D, F) in forearm, buttock and cheek for 154 subjects. Red and blue indicate positive and negative Spearman’s rho values, respectively.

**Fig. 3.**
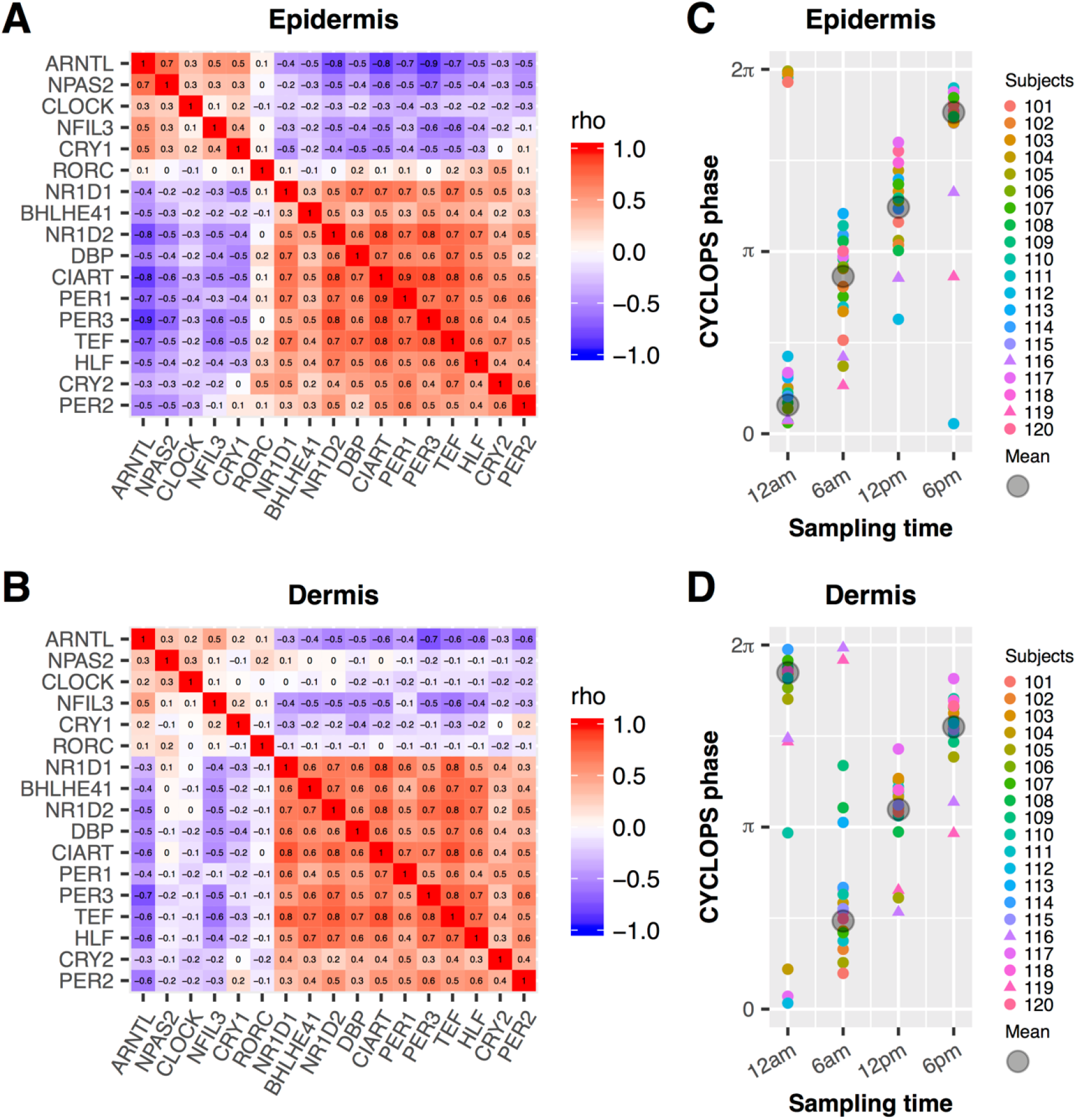
Evaluation of clock function and accuracy of CYCLOPS ordering in epidermis and dermis. A heatmap of Spearman’s rho for clock and clock-associated genes from longitudinal (n= 20, Additional file 1: Table S1) and population-based data (n = 154, Additional file 1: Table S1) is shown for epidermis (A) and dermis (B). The population-based epidermis or dermis samples are from forearm, cheek and buttock of the same 154 subjects. The longitudinal order of samples collected from 20 subjects were accurately recalled in epidermis (C) and dermis (D). Different colors indicate different subjects, and the circular average phases for all samples is shown in grey at each sampling time. Samples from subjects 116 and 119 are indicated by triangles.

### Genome-wide rhythms in epidermis and dermis across body sites

Based on the evidence for functional clocks in epidermis and dermis, we explored their rhythmic transcriptomes. Because biopsy time was not recorded for any of the population samples, we used CYCLOPS to reconstruct their temporal order, as reported previously [18]. Samples from epidermis and dermis were ordered separately. To maximize the oscillation signal, we added a random sampling step (described in Methods). Using this strategy, we achieved high-quality ordering with 97% (epidermis) and 95% (dermis) samples. Assessment of quality was based on two primary criteria: (1) CYCLOPS-predicted sampling times for the 20 subjects with longitudinal data matched *known* sampling times (Fisher’s circular correlation of 0.973 in epidermis and 0.925 in dermis) (Fig. 3C-D), and (2) the predicted phases of clock genes matched the mouse reference (Additional file 1: Fig. S2A-B, outer versus inner circle; Fisher’s circular correlation, 0.74 in epidermis and 0.71 in dermis).

We then analyzed 24 h patterns of expression for all transcripts genome-wide as a function of predicted sampling times. 171 and 70 genes met our criteria for rhythmicity in epidermis and dermis, respectively (modified cosinor regression, *P* < 0.01, relative amplitude (rAMP) > 0.1, goodness-of-fit (rsq) > 0.1, and fitmean > 16) (Fig. 4A and Additional file 3). 41 (24%) of these genes from epidermis and 30 (43%) genes from dermis had robustly cycling homologs in mouse telogen, suggesting strong evolutionary conservation. Mouse telogen refers to the non-proliferative stage in skin, which has stronger circadian output than the hair follicle growth stage [25]. Most of these conserved genes (Fig. 4B and C) oscillated with similar phases (relative to *ARNTL*/*Arntl* phase) between species (within 3 h difference). Eighteen of them were rhythmically expressed in *both* epidermis and dermis, 12 of which were clock and clock-associated genes. Interestingly, 4 of the 6 remaining genes (*WEE1*, *TSC22D3*, *FKBP5* and *KLF9*), although not currently considered clock-associated, were circadian-expressed in many mouse tissues (JTK Q-value < 0.05 in 7-13 mouse tissues from CircaDB) [26].

**Fig. 4.**
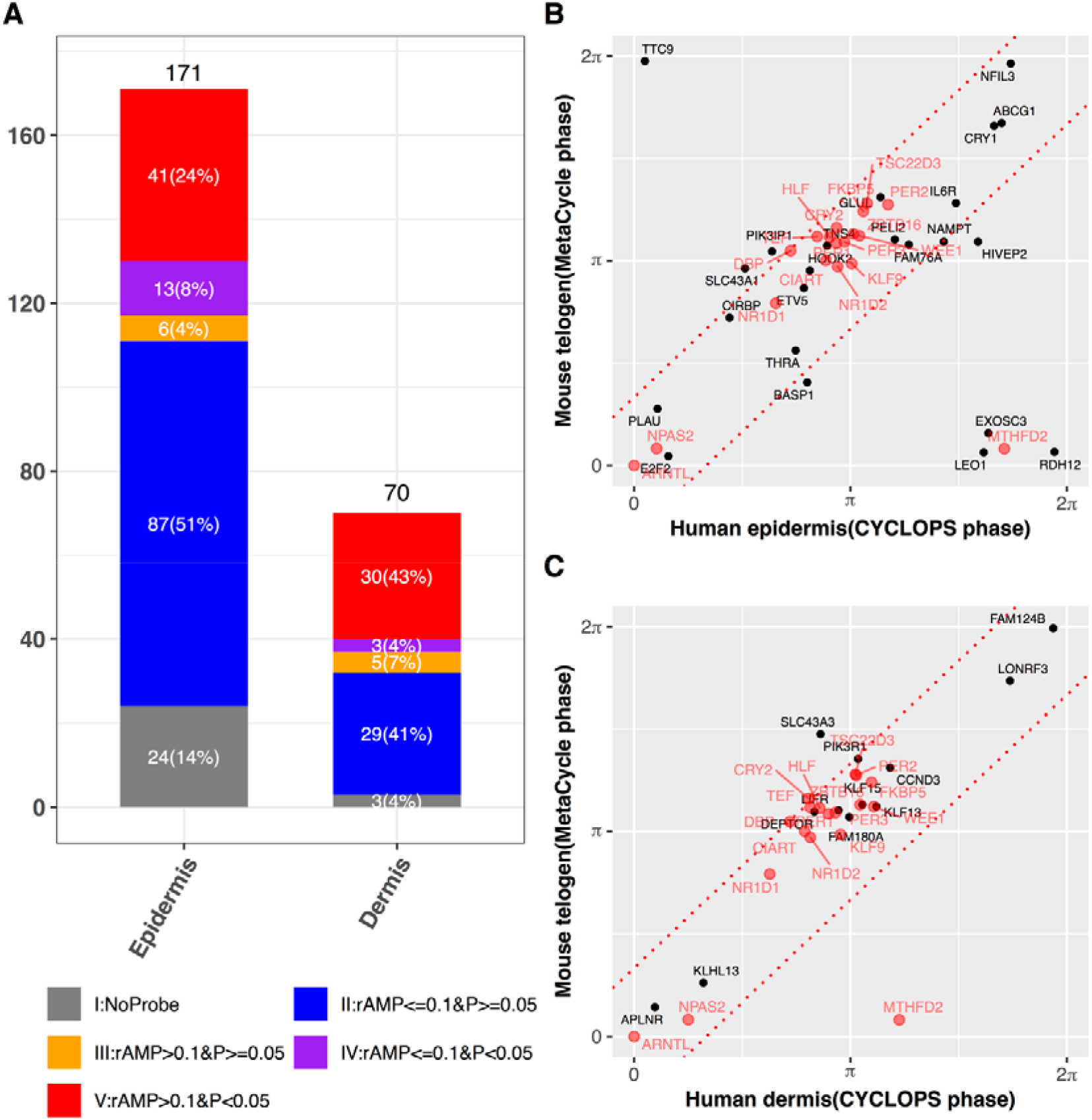
Conserved circadian transcriptional output between human and mouse skin. (A) Circadian genes in human epidermis and dermis were grouped into 5 categories: (I) no mouse homolog (grey), (II) low amplitude (rAMP <= 0.1) and not cycling (*P* > 0.05) (blue), (III) high amplitude (rAMP > 0.1) and not cycling (orange), (IV) low amplitude and cycling (*P* < 0.05) (purple), (V) high amplitude and cycling (red). (B and C) Phase comparison of high amplitude cycling genes (Group V, n = 41 for epidermis and n = 30 for dermis) reveals strong phase conservation between mice and humans. Overlapped circadian genes in epidermis and dermis are shown in red. The gene phases were adjusted to *ARNTL/Arntl* phase (set to 0). The red dotted line indicate the phase difference with □/4. Mouse data are from Geyfman et al. [25] (Additional file 1: Table S1).

### The epidermal layer provides the better marker of circadian phase

To evaluate which skin layer is the more reliable predictor of phase, we used ZeitZeiger [20] to identify candidate biomarker sets. ZeitZeiger was trained by a combination of population and longitudinal samples. There were two separate training datasets for CYCLOPS-ordered epidermis (n=483) and dermis (n=472) samples from forearm, buttock and cheek. Training produced candidate biomarker sets of 12 and 21 genes for epidermis (Fig. 5A) and dermis (Additional file 1: Fig. S3A), respectively.

**Fig. 5.**
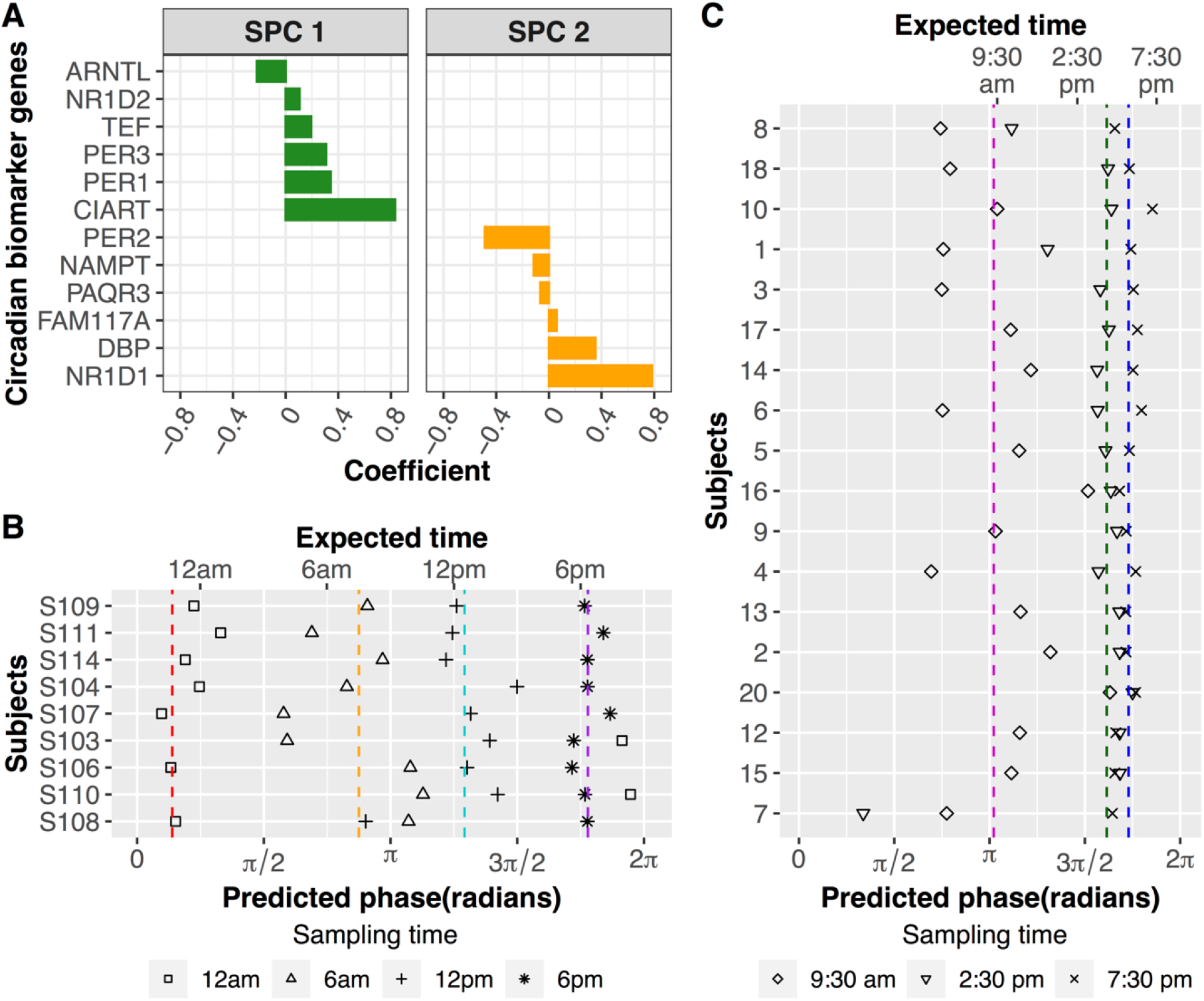
Population-level circadian biomarkers work for a single human epidermis sample. (A) 12 circadian marker genes in the first two sparse principal components (SPCs) were selected by ZeitZeiger. (B) Validation of circadian biomarkers using epidermis samples from 9 subjects that were excluded from the training set. The epidermis samples were collected every 6 h over a circadian day. Average predicted phases of samples collected at 12 AM, 6 AM, 12 PM and 6 PM are indicated with red, orange, cyan and purple dashed lines, respectively. (C) Validation of circadian biomarkers using longitudinal epidermis samples from 18 subjects in Sporl F. et al. [27] (Additional file 1: Table S1). The epidermis samples were collected every 5 h from 9:30 AM to 7:30 PM. Average predicted phases of samples collected at 9:30 AM, 2:30 PM and 7:30 PM are indicated with magenta, green and blue dashed lines, respectively.

We tested the biomarker sets in 36 epidermis and dermis samples from the same 9 subjects. Each of these subjects contributed 4 time-stamped samples that were not part of the training set. Whereas the epidermis set reproduced known sampling time in 8/9 subjects, the dermis set only performed in 5/9 (Fig. 5B and Additional file 1: Fig. S3B). The average predicted phase was within 3 h of the expected sampling time for 4/4 (epidermis) and 3/4 (dermis) time points. Overall, the average absolute error across different times of day for the 9 subjects was 2.5 h for epidermis compared to 3.8 h for dermis (Additional file 1: Fig. S4A and S4B). In summary, the epidermis was the better source of biomarkers of circadian phase.

### A circadian biomarker that is robust to a different experimental design

But how well does this biomarker perform with different sample collection methods and experimental designs? We tested its prediction accuracy in an independent dataset obtained from suction-blister epidermal samples collected over 24 h from 18 Caucasian subjects in Germany [27]. The epidermal biomarker reproduced the time-stamped sample order in 14 of 18 subjects (Fig. 5C). One sample phase in each of the four remaining individuals (subject 20, 12, 15, and 7) were poorly predicted. The overall average absolute error was 3.1 h. For future validation of this biomarker in diverse populations, diseases, and experimental designs, we developed SkinPhaser (https://github.com/gangwug/SkinPhaser) to enable the biomarker test in public datasets.

## Discussion

We previously reported 29 genes from epidermal forearm skin whose expression values could accurately determine circadian phase to within 3 h [18]. The work here improves upon this and brings us closer to a clinically-viable biomarker of body time.

First, a biomarker should be stable. We demonstrate this across multiple different body sites, sun- and non-sun-exposed. Second, a biomarker should be practical; its value must exceed its cost. We demonstrate a smaller set (12 genes) with improved accuracy from a single biopsy of the skin’s most superficial layer. Third, an ideal biomarker performs regardless of detection platform. We demonstrate this for two different array platforms (Agilent vs. Affymetrix) and collection strategies (suction blister vs. punch biopsy). Finally, a circadian biomarker should generalize to a diverse population; circadian medicine may apply to everyone. We demonstrate this for different geographic groups (Germany and U.S.). Nevertheless, testing of circadian biomarkers on a broader range of datasets is critical. To help drive this forward, we developed SkinPhaser – a user-friendly app to benchmark this biomarker set across subject demographics, disease states, and experimental designs.

In both longitudinal and population datasets, clock gene phases were similar between epidermis and dermis. This suggested that circadian phase is coherent across skin layers. Interestingly however, the robustness of molecular oscillations differed between the two layers. Why is the clock in epidermis stronger than dermis? There are several possibilities. The dermal layer is much larger and more heterogeneous. Some cell types in the dermis may have weak intrinsic oscillators. Another possibility is that epidermal clocks may entrain more efficiently by virtue of their direct contact with environmental cues. Finally, systemic cues in the circulation may dampen dermal oscillations as the dermis is perfused, whereas the epidermis is not [28].

Future tests need to link biomarker predicted phase to different physiological measurements and confirm the test consistency across a range of disease states and pathologies. Clinical use of circadian biomarkers will require fast, cost-effective, and non-invasive sampling techniques and an optimal detection platform. Finally, head-to-head comparisons are necessary to link the molecular clock phase predicted from skin biomarkers with the melatonin phase predicted from blood [11,13–15,17] and other physiological phases (e.g., core body temperature) [29].

With continuing efforts from multiple groups in the last 30 years, the field is moving closer to the clinical version of circadian biomarkers. Possible applications include tailoring medication dosage times, deciding on procedural timing, and assisting in the clinical diagnosis and treatment of circadian rhythm sleep disorders. These future clinical tests on circadian biomarkers will make medicine more precise.

## Conclusions

Our study shows that a functional clock running in human epidermis (12 biomarker genes) is superior to dermis (21 biomarker genes) to report circadian phase regardless of body site. Using one epidermis sample, this biomarker set performs well across body sites, ages, sexes, and detection platforms. Finally, we developed SkinPhaser – a user-friendly app to use these biomarkers on community datasets. Future studies should focus on developing a minimally invasive procedure for obtaining intact epidermal RNA. Overcoming these roadblocks will enable widespread adoption of circadian medicine.

## Methods

### The collection of human longitudinal dermis samples

20 healthy Caucasian male subjects from the U.S. were recruited for the longitudinal dermis sample collection. These were the same 20 subjects in our previous collection of longitudinal epidermis samples [18]. Each subject was provided informed consent. Associated protocol was approved by an Institutional Review Board (Aspire; http://aspire-irb.com/). Subjects donated one skin punch biopsy at each of four time points (6 AM, 12 PM, 6 PM, and 12 AM) over a 24-h period. Biopsies were separated into epidermal and dermal layers by laser capture microdissection (LCM). The mRNA extraction, target labeling and hybridization to microarrays were described previously [23].

### The MetaCycle analysis of human longitudinal dermis samples

The RMA algorithm from affy R package [30] was used to extract the expression profile from the raw CEL files of 79 human longitudinal dermis samples (Additional file 1: Table S1). The expression profile was analyzed with the meta3d function of MetaCycle R package [21] with default settings, except ‘cycMethodOne’ = “ARS”, ‘minper’ = 24, and ‘maxper’ = 24. ARSER [31] was used to analyze the time-series data individual by individual. Then meta3d was used to integrate the analysis results from 19 subjects (subject 115 had one missing time point and was excluded from the longitudinal analysis). Circadian genes were defined by *P* < 0.05 and rAMP > 0.1. We applied a less strict cut-off (*P* < 0.1 and rAMP > 0.1) for evaluating overlap between human epidermis and dermis and phase set enrichment analysis (PSEA) [22].

### Comparing clock robustness between human epidermis and dermis

We used the microarray dataset from a previous population study of human epidermis and dermis [23]. This study recruited 154 Caucasian females (aged 20-74 y) in the U.S. Three punch biopsies were collected from each subject, with one biopsy per body site (arm, buttock and cheek). Sample collection times were not recorded. Epidermal and dermal layers were separated by LCM. This yielded six groups of skin samples: arm epidermis, buttock epidermis, cheek epidermis, arm dermis, buttock dermis and cheek dermis. We measured pairwise clock gene correlations [24] between epidermis and dermis at each body site.

### The hybrid design and seed circadian gene lists

We used the hybrid design to order epidermis and dermis samples across body sites [18]. In detail, the RMA algorithm was applied to all 533 epidermis samples (79 longitudinal samples from arm, and all population-based samples from arm, buttock and cheek except 8 missing samples). ComBat [32] corrected for batch effects. Probe sets were annotated with gene symbols, and one representative probe set with the maximum median absolute deviation was selected for each gene. The same steps were performed for 531 dermis samples (79 longitudinal samples from arm, and all population-based samples from arm, buttock and cheek except 10 missing samples).

### Ordering human epidermal and dermal samples using revised CYCLOPS pipeline

The revised CYCLOPS pipeline [18] was used to order human epidermal and dermal samples. Prior to ordering, we added a down-sampling step [4] to enrich for samples with maximum clock oscillation signal. Maximum clock oscillation signal was defined as the highest Z-statistic (Mantel test) compared to the benchmark correlation matrix (Additional file 5) of 298 previously ordered human epidermis samples [18]. In sum, 519 (~97%) of epidermis samples and 506 (~95%) of dermis samples were selected for CYCLOPS ordering.

To further optimize CYCLOPS ordering quality, tested three different seed lists as input to the algorithm: 1) 158 genes that are identified as circadian genes in at least two of four longitudinal skin datasets: human dermis from this study, human epidermis [18], and mouse time-series telogen and anagen [25]; 2) human homologs of genes cycling in at least 9 of 12 mouse tissues [3]; 3) 17 clock and clock-associated genes. See Additional file 4. Assessment of ordering quality was based on two fisher circular correlations values: 1) sampling time correlation value and 2) clock gene correlation value (Additional file 1: Fig. S5). For epidermis, best results were obtained from 519 samples ordered using seed set #3 (above). For dermis, best results were obtained from 506 samples ordered using seed set #2. Circadian genes were selected with *P* < 0.01, fitmean > 16, rAMP > 0.1, and rsq > 0.1 from the cosinor regression analysis results (Additional file 3).

### ZeitZeiger to identify biomarkers of circadian phase

ZeitZeiger was used to identify circadian biomarkers from CYCLOPS-ordered human epidermis and dermis samples. Epidermis or dermis samples were divided into two groups: testing and training. The testing set in epidermis (and dermis) included 36 time-stamped samples from nine subjects (each subject with four samples). The epidermis (and dermis) training set included all ordered samples from the population group and from the remaining 11 subjects of longitudinal group. In total, 483 epidermis samples and 472 dermis samples were used for training ZeitZeiger. Default ZeitZeiger settings were used for training samples, with two modifications: normalizing the expression value of each gene and selecting dynamic genes. There are three steps in expression normalization: 1) normalizing by three non-circadian genes in skin (*GPKOW*, *BMS1* and *ANKFY1*), 2) rounding the expression outliers, and 3) normalizing the expression value of each gene using its maximum expression value across all samples. The dynamic genes were selected with an rsq value > 0.1 from cosinor regression analysis of the optimal CYCLOPS ordering of epidermis and dermis samples. We ran ZeitZeiger using sumabsv = 2 and sumabsv = 3 in epidermis and dermis, respectively. Those genes in the first two SPCs with an absolute coefficient value > 0.05 were selected as candidate biomarkers of circadian phase.

### Validation of circadian phase prediction across different experiment designs

The prediction accuracy of 12 epidermal or 21 dermal genes was further evaluated using the 36 time-stamped epidermis or dermis samples in the testing set (Additional file 1: Figure S6). The phase difference between any two time-stamped samples for the same subject is known and is not influenced by the individual chronotype. The prediction accuracy is thus evaluated by comparing the difference between predicted and known sample phases.

The epidermis biomarker genes were then tested in an independent longitudinal dataset [27] of 20 Caucasian male and female subjects in Germany. Epidermis samples were harvested at 9:30 am, 2:30 pm and 7:30 pm using suction blister method, with transcriptome detection on an Agilent platform. Those 18 subjects with 3 epidermis samples were used for testing the prediction accuracy of 12 epidermal circadian marker genes (Additional file 1: Figure S6).

## Abbreviations

CYCLOPS: cyclic ordering by periodic structure
DLMO: dim-light melatonin-onset
GPCR: G protein-coupled receptor
LCM: laser capture microdissection
PSEA: phase set enrichment analysis

## Declarations

### Ethics approval and consent to participate

Associated protocols were approved by Institutional Review Board (http://aspire-irb.com/).

### Consent for publication

Not applicable.

### Availability of data and materials

The gene expression data of longitudinal human dermis have been deposited in GEO (GSE139300). The GEO accession numbers for other published skin datasets used in this study were listed in Additional file 1: Table S1. The versions of software packages and their available links were listed in Additional file 1: Table S2.

### Author contributions

J.B.H., J.E.O., and K.J.M, designed research; G.W., M. D. R, J.D.S. and L. J. F. performed research; G.W., J. B. H., and M. D. R. contributed analytic tools; J. B. H., G. W., M. D. R., L. J. F., D. F. S. and J.E.O., and K.J.M wrote the paper. All co-authors contributed to and approved the final draft.

## Acknowledgments

We thank all the members of Hogenesch lab for thoughtful discussions.

## Supplementary material

Additional file 1: supplemental figures and tables (docx)

Additional file 2: the reference correlation matrix of 17 mouse clock and clock associated genes (csv)

Additional file 3: circadian genes identified from human epidermis and dermis (xlsx) Additional file 4: seed gene lists used for CYCLOPS ordering (csv)

Additional file 5: the benchmark correlation matrix of 298 previously ordered human epidermis samples (csv)

