## Additional file 1 for "A single-sample circadian biomarker that performs across populations and platforms"

Fig. S1. Comparison of circadian genes identified from time-series analyses of human dermis and epidermis

Fig. S2. Phase order of identified clock and clock-associated genes from human epidermis and dermis

Fig. S3. Biomarker candidate genes for predicting circadian phase of a single human dermis sample

Fig. S4. Prediction accuracy of circadian biomarkers from epidermis and dermis

Fig. S5. The pipeline of CYCLOPS ordering epidermis and dermis samples

Fig. S6. The pipeline of identifying and testing epidermal circadian biomarkers using ZeitZeiger

Table S1. The list of skin datasets used in this study

Table S2. The list of software packages used in this study

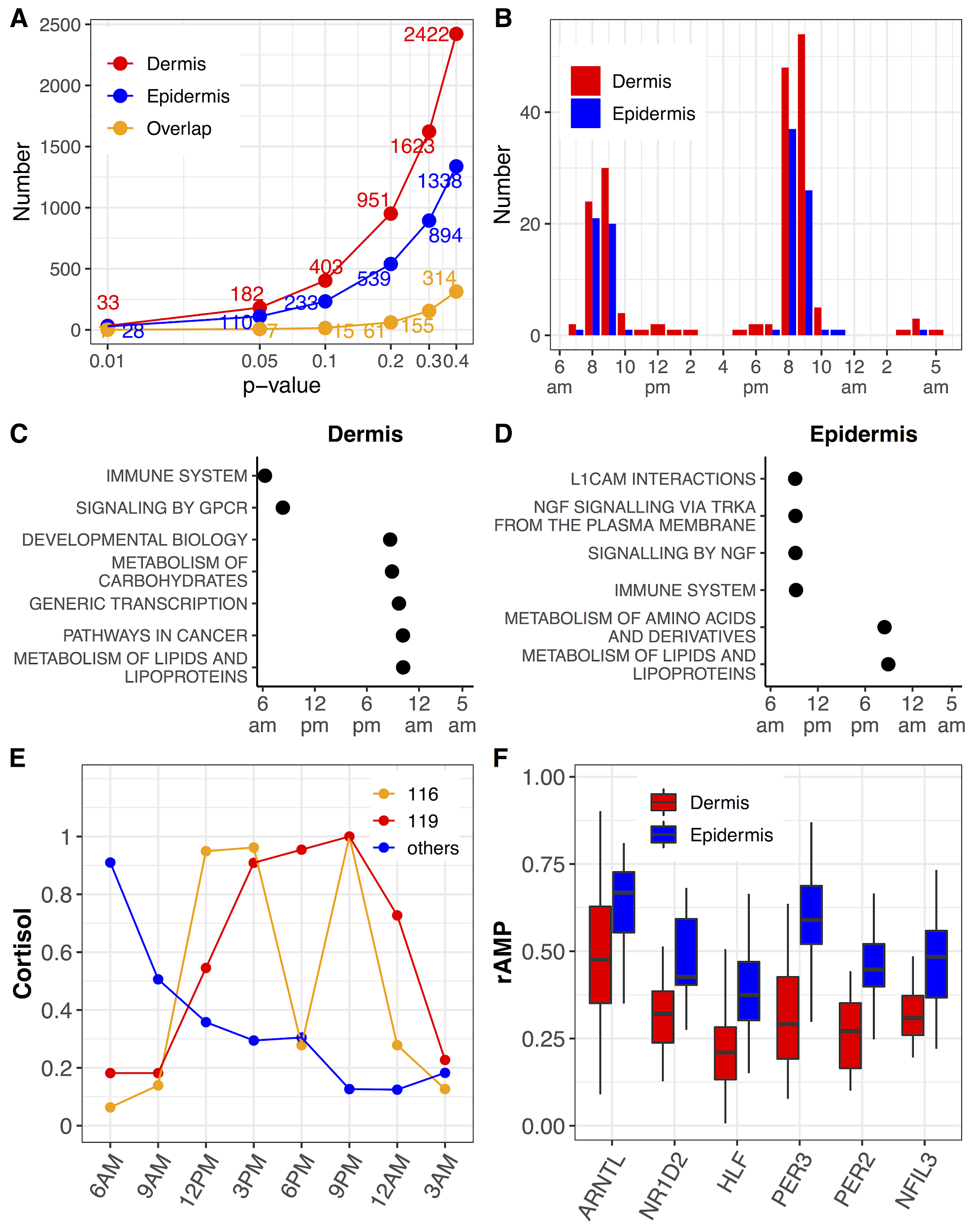

Fig. S1. Comparison of circadian genes identified from time-series analyses of human dermis and epidermis. (A) Number of circadian genes at a series of *P* value cut-offs were shown in dermis and epidermis. (B) Phase distribution of circadian genes (*P* < 0.05) identified in epidermis and dermis. (C and D) Significant enriched time-dependent pathways of circadian genes identified in dermis and epidermis. A less significant cut-off (*P* < 0.1) was used to select circadian genes for performing PSEA analysis. (E) Salivary cortisol levels in subjects 116 and 119 along time, comparing to the average cortisol levels of other subjects along time. The salivary cortisol data is from our previous study [[18]](https://paperpile.com/c/bc07Ux/EExS). The y-axis indicates the relative cortisol level normalized by maximum cortisol value. (F) The individual rAMP of six clock genes were shown for dermis and epidermis.

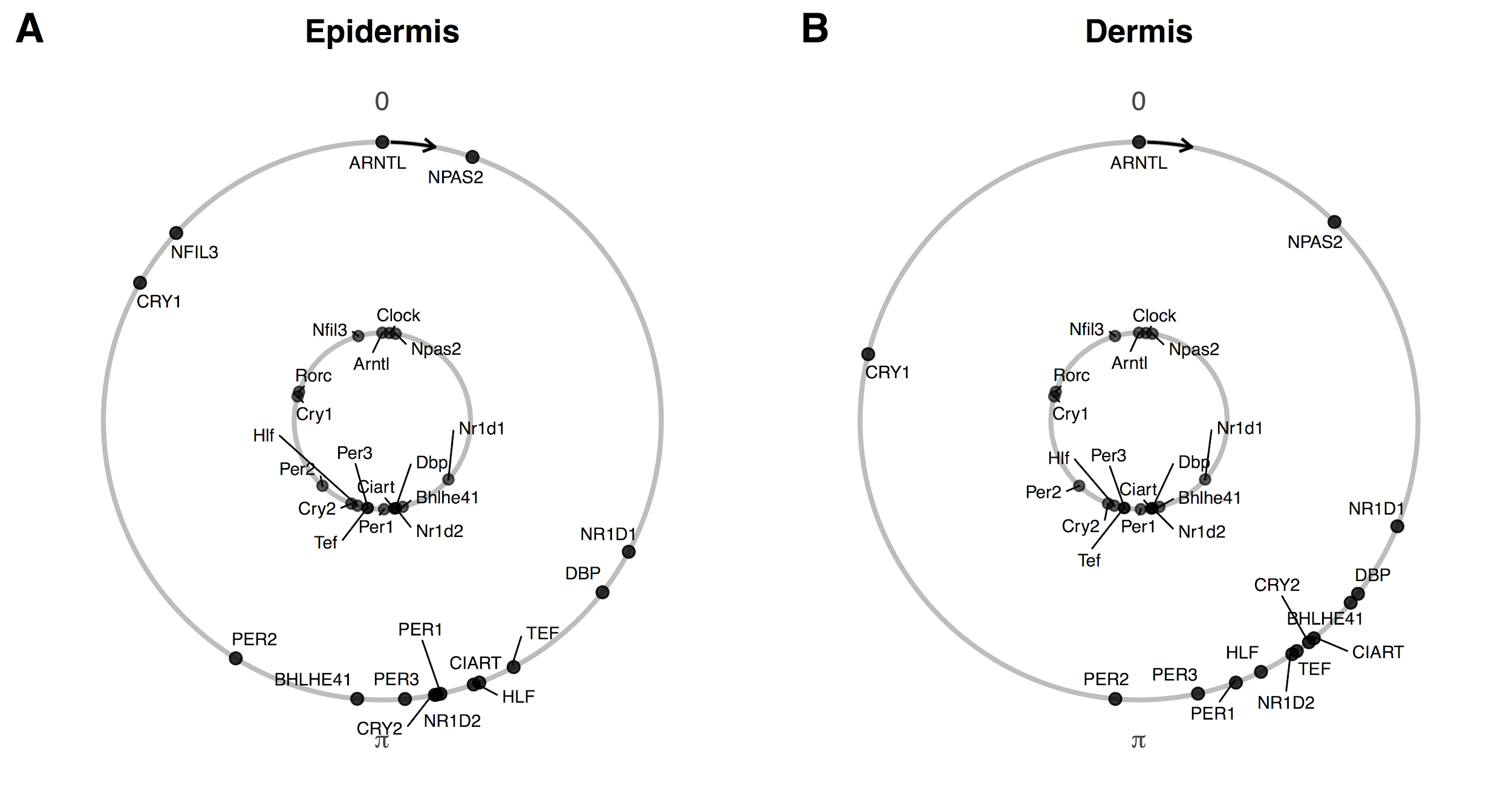

Fig. S2. Phase order of identified clock and clock-associated genes from human epidermis and dermis. Conserved phase relationships are shown for clock and clock-associated genes (internal circle, mouse; external circle, human) in human epidermis (A) and dermis (B). The phase of *ARNTL/Arntl* is set as 0.

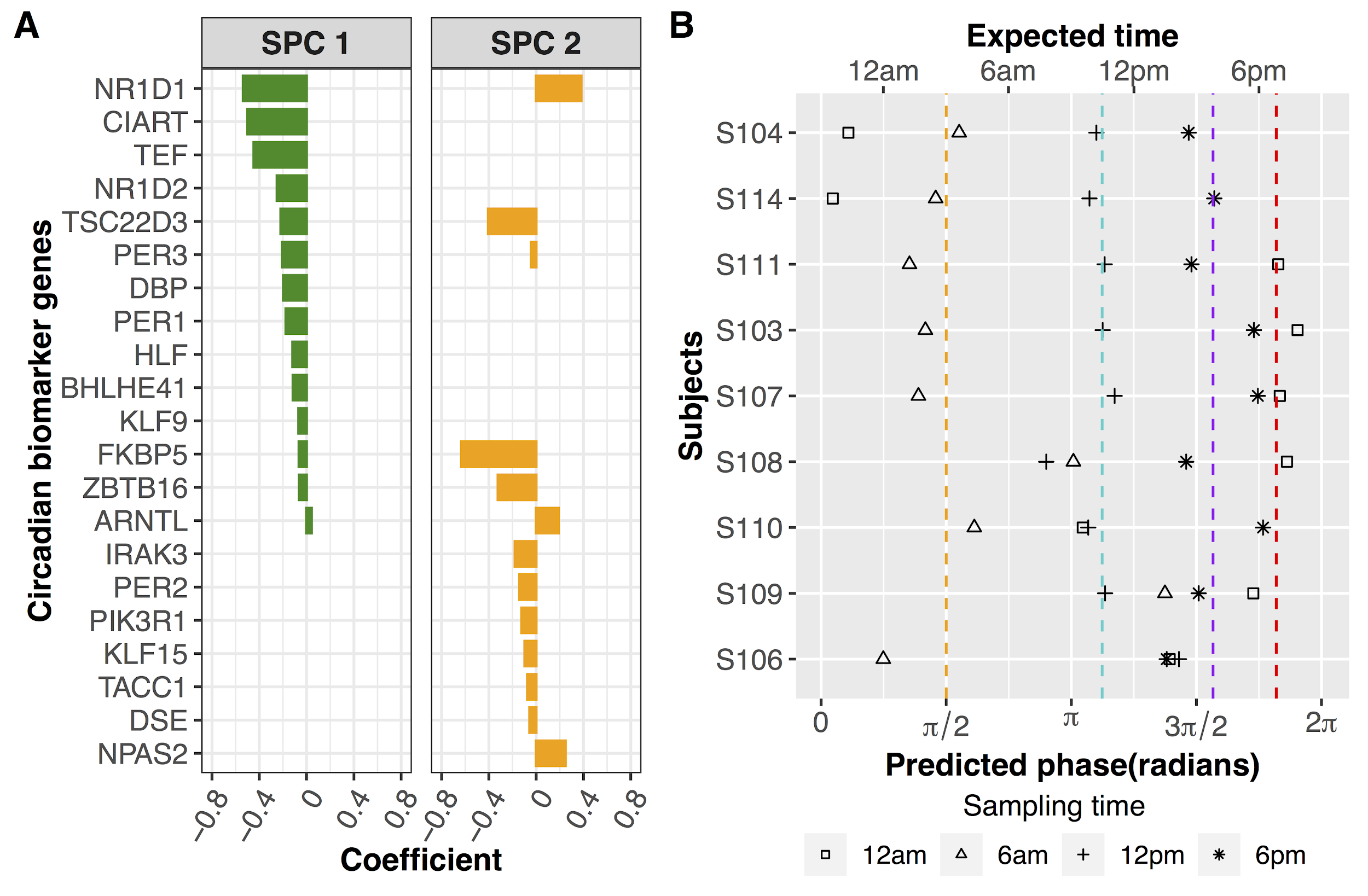

Fig. S3. Biomarker candidate genes for predicting circadian phase of a single human dermis sample. (A) 21 candidate biomarker genes were selected by ZeitZeiger. (B) Validation of predicted markers using dermis samples from 9 subjects that were excluded from the training set. The dermis samples were collected every 6 h over a circadian day. Average predicted phases of samples collected at 12 AM, 6 AM, 12 PM and 6 PM are indicated with red, orange, cyan and purple dashed lines, respectively.

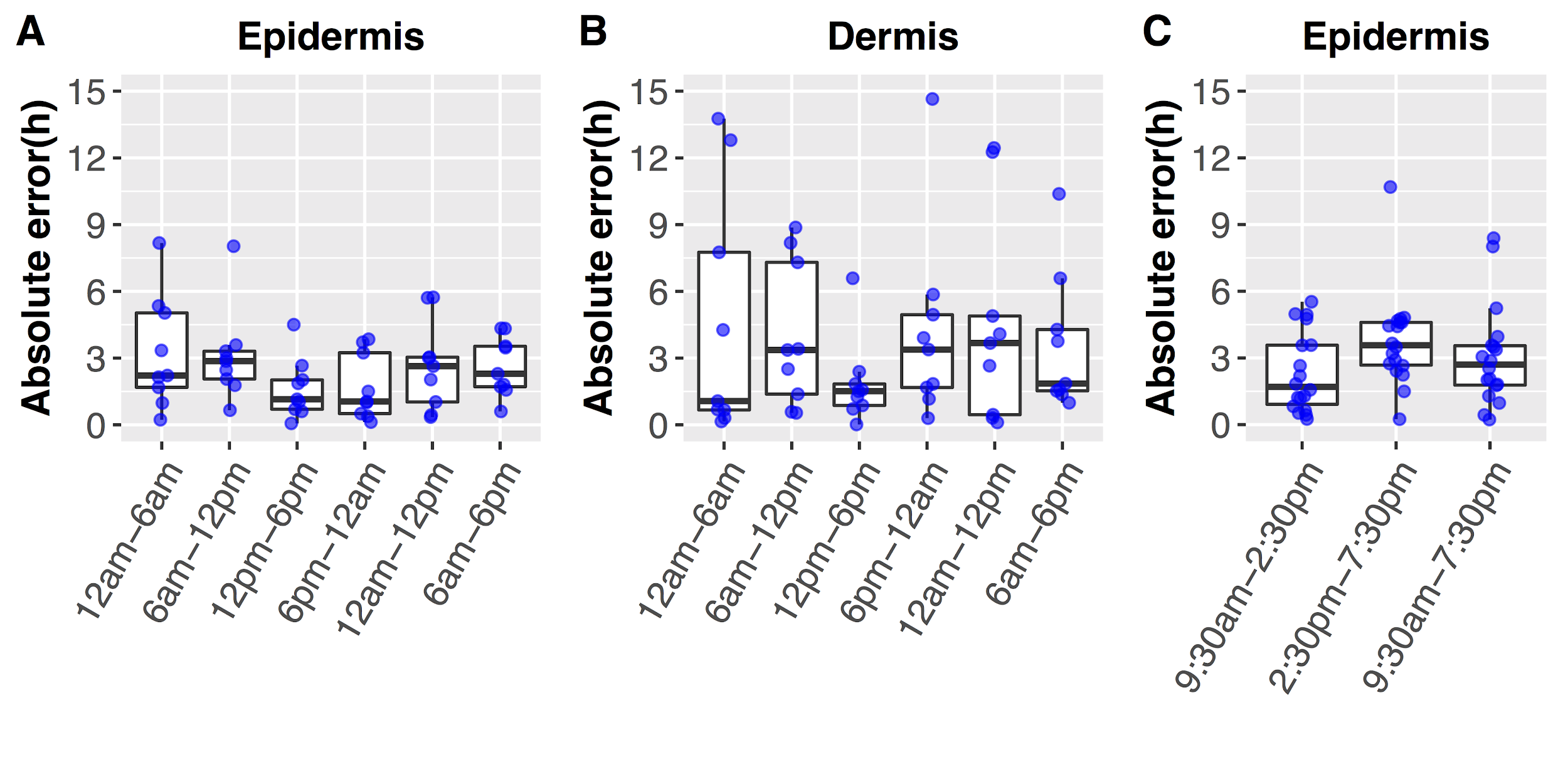

Fig. S4. Prediction accuracy of circadian biomarkers from epidermis and dermis. Absolute errors of phase prediction using selected biomarker genes are plotted for nine subjects at six time windows for epidermis (A) and dermis (B). Absolute errors of phase prediction using biomarker genes from epidermis are plotted for 18 subjects at three time windows (C) using data from Sporl et al. study [[27]](https://paperpile.com/c/bc07Ux/A4Dw) (Table S1).

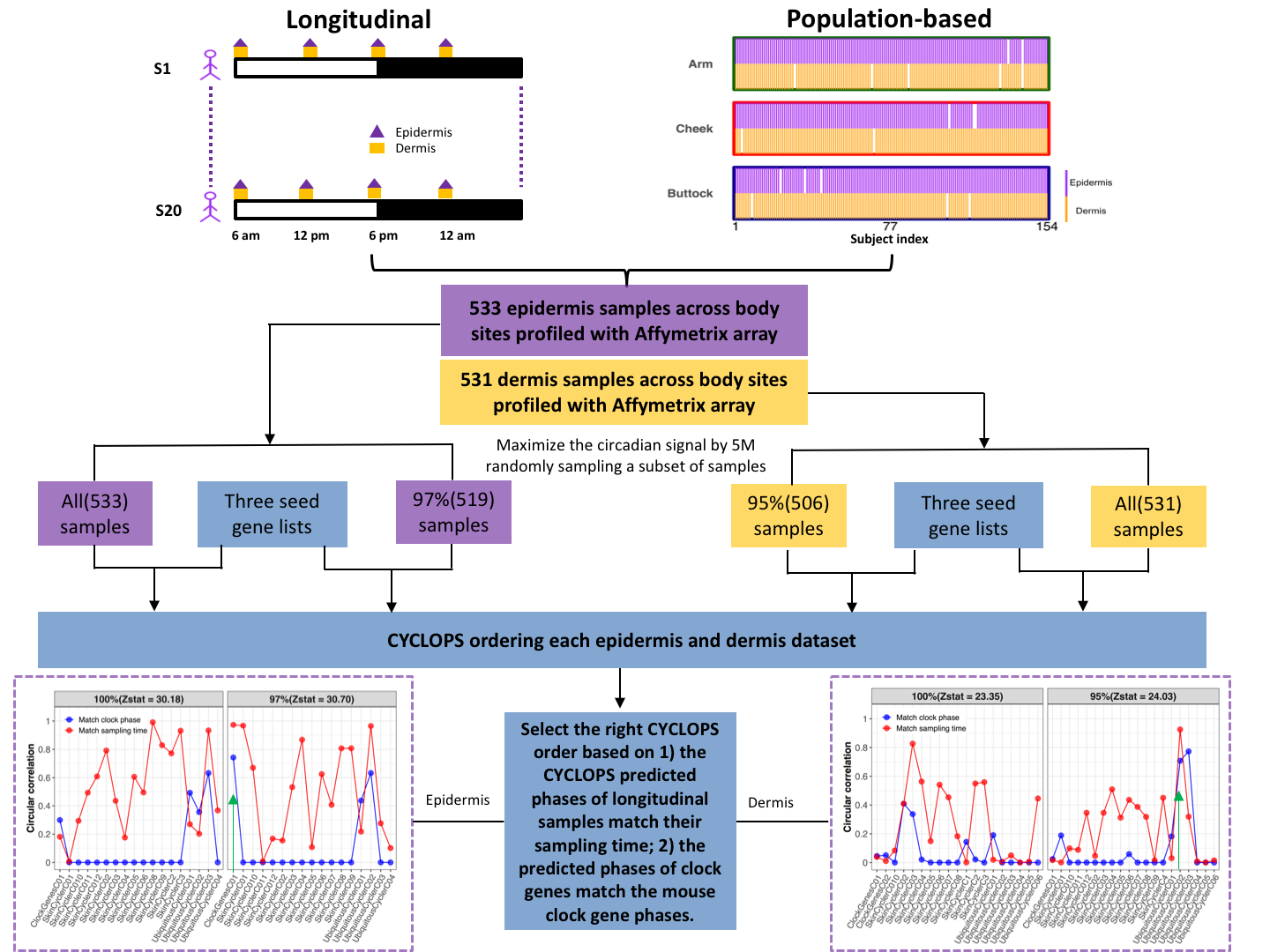

Fig. S5. The pipeline of CYCLOPS ordering epidermis and dermis samples. We got 533 epidermis samples and 531 dermis samples by combining longitudinal samples collected from 20 subjects and population-based samples collected from 154 subjects at three body sites (arm, cheek and buttock). We also got 97% (519) and 95% (506) of epidermis and dermis samples through 5 million randomly sampling for maximizing the circadian signals in the selected subset of epidermis and dermis samples. Thus each skin layer has two datasets. Three seed circadian gene lists were used for CYCLOPS ordering these four datasets, which include 158 skin circadian genes, human homologs of mouse ubiquitous circadian genes, and 17 clock and clock-associated genes (Additional file 4). The optimal CYCLOPS order should meet two criteria: 1) the CYCLOPS predicted phases of longitudinal samples match their known sampling time (match sampling time); 2) the phase of clock genes from ordered human epidermis or dermis samples meet with the phase order of mouse clock genes (match clock phase). The matchable of sampling time and clock phase are reflected by the corresponding circular correlation value. If fewer than 8 cycling clock genes were identified (*P* < 0.01, fitmean > 16, rAMP > 0.1, and rsq > 0.1), the circular correlation value for ‘match clock phase’ was set as 0. The optimal CYCLOPS order was indicated with green arrow, with both high circular correlation values of sampling time and clock phase.

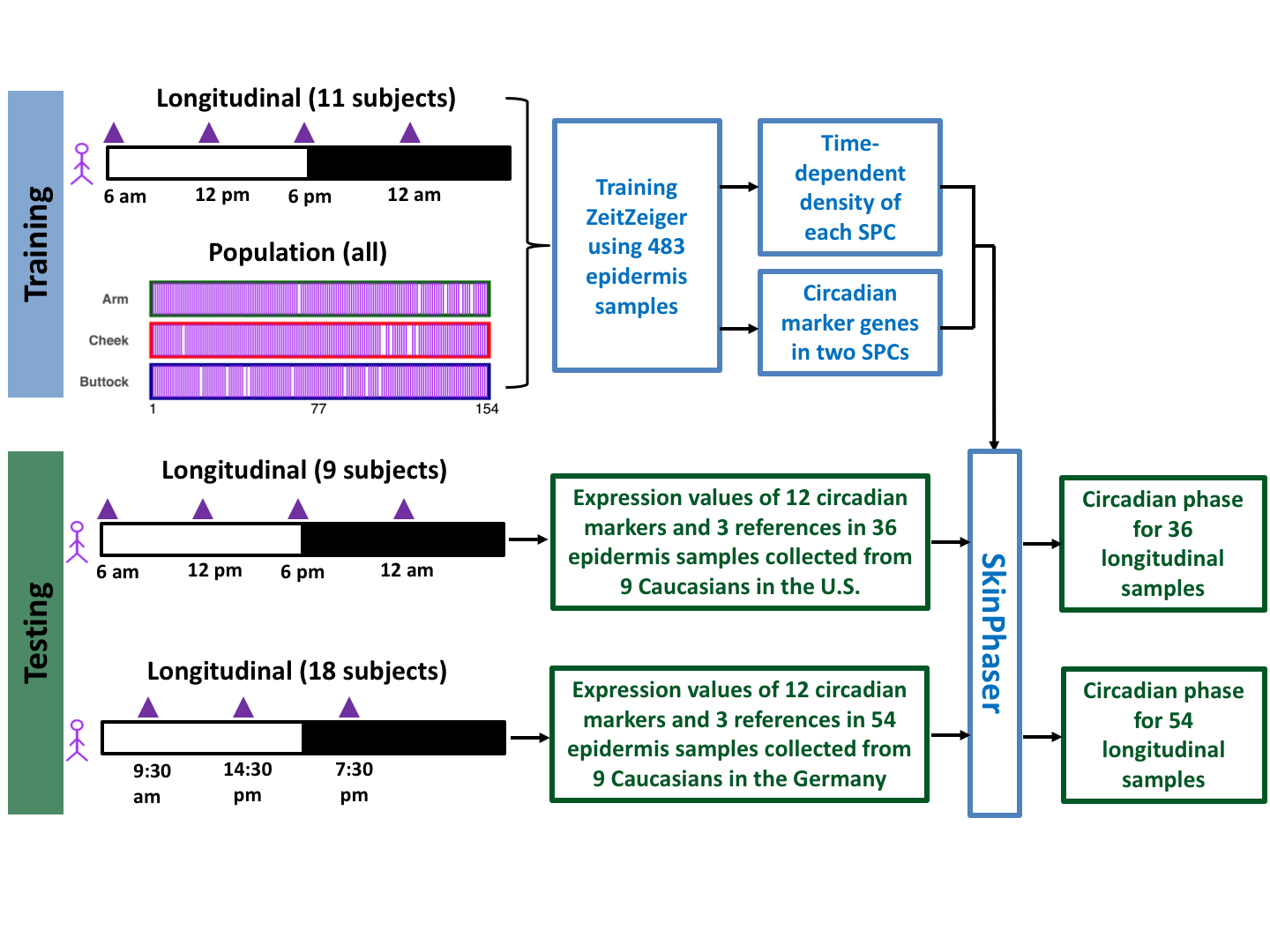

Fig. S6. The pipeline of identifying and testing epidermal circadian biomarkers using ZeitZeiger. The training dataset includes 483 longitudinal and population-based epidermis samples. Using this training dataset, a set of 12 circadian biomarker genes were selected by ZeitZeiger. The user-friendly SkinPhaser app was developed for predicting circadian phase using this set of 12 circadian biomarker genes. The prediction accuracy of circadian biomarkers was further tested using two testing datasets. One dataset includes 36 epidermis samples collected from 9 male subjects in the U.S., and each subject donated one sample at each of four time points (6 AM, 12 PM, 6 PM and 12 AM). The other one includes 54 epidermis samples collected from 18 male and female subjects in Germany, and each subject donated one epidermis sample at each of three time points (9:30 AM, 14:30 PM and 7:30 PM).

**Table S1. The list of skin datasets used in this study**

| **Datasets** | **Reference** | **Platform** | **Site** | **#Samples** | **Experimental design** |
| --- | --- | --- | --- | --- | --- |
| 1. Mouse telogen  (longitudinal) | Geyfman M., et al., 2012, PNAS. GSE38622 | MoGene-1_0-st | telogen | 13 | C57BL/6CR mice were housed under 12 h:12 h LD cycles with food and water ad libitum. Whole skin was collected at 4-h intervals for 48 h. Telogen samples were collected from P46 mice. Equal amounts of RNA from the three mice for each time point were pooled. |
| 2. Epidermal  skin from Caucasian in Germany  (longitudinal) | Sporl F, et al., 2012, PNAS. GSE35635 | Agilent- 014850 | NA | 54 | Suction blisters for 10 male and 10 female volunteers aged 28.35 ± 4.36 y (mean ± SD) were harvested at 9.30 am, 2.30 pm and 7.30 pm. Those 18 subjects with 3 samples were selected in this study. |
| 3. Epidermal  skin from Caucasian in the U.S.  (longitudinal) | Wu G., et al., 2018, PNAS.  GSE112660 | HG-219 array | forearm | 79 | Four forearm skin samples were collected at 12am, 6am, 6pm, 12pm for each of 20 male subjects, except one missing sample from subject 115. The ages of these 20 subjects are between 21 and 49 years old. LCM was performed to separate dermis from epidermis. |
| 4. Dermal  skin from Caucasian in the U.S.  (longitudinal) | This study.  GSE139300 | HG-219 array | forearm | 79 | As described above for dataset 3. |
| 5. Epidermal skin from Caucasian in the U.S. (population) | Kimball A. B., et al., 2018, J Am Acad Dermatol.  GSE112660, GSE139305 | HG-219 array | forearm,  cheek and buttock | 454 | One skin sample from forearm, cheek and buttock was taken from each of 154 female subjects, aged between 20 and 74 years old. Samples were designed to collect during the working hours, between 9am to 5pm. LCM was performed to separate dermis from epidermis. There are 17 subjects with one missing sample and one subject with two missing samples. |
| 6. Dermal skin from Caucasian in the U.S. (population) | Kimball A. B., et al., 2018, J Am Acad Dermatol.  GSE112660, GSE139305 | HG-219 array | forearm,  cheek and buttock | 452 | As described above for dataset 6. |

**Table S2. The list of software packages used in this study**

| **Software package** | **Version** | **Link** |
| --- | --- | --- |
| affy | 1.62.0 | <https://bioconductor.org/packages/release/bioc/html/affy.html> |
| ape | 5.3 | <https://cran.r-project.org/web/packages/ape/index.html> |
| CircStats | 0.2-6 | <https://cran.r-project.org/web/packages/CircStats/index.html> |
| CYCLOPS | v3.0.2.1 | <https://github.com/gangwug/CYCLOPSv3.0.2.1> |
| julia | 0.3.12 | <https://julialang.org/downloads/oldreleases.html> |
| limma | 3.40.2 | <https://bioconductor.org/packages/3.9/bioc/src/contrib/Archive/limma/> |
| MetaCycle | 1.2.0 | <https://cran.r-project.org/web/packages/MetaCycle/index.html> |
| Oscope | 1.14.0 | <http://bioconductor.org/packages/release/bioc/html/Oscope.html> |
| PSEA | 1.1 | <https://github.com/ranafi/PSEA> |
| R | 3.6.1 | <https://www.r-project.org/> |
| sva | 3.32.1 | <https://bioconductor.org/packages/release/bioc/html/sva.html> |
| zeitzeiger | 2.0.1 | <https://github.com/hugheylab/zeitzeiger> |
